# Relative expression of *THY1* and *PDGFRA* in equine endometrosis in relation to severity, histopathological features and endometrial transcripts

**DOI:** 10.64898/2026.11.07.764916

**Authors:** Łukasz Zdrojkowski, Anna Niwińska, Ewa Kautz-Wasilewska, Dawid Tobolski, Magdalena Fajkowska, Małgorzata Rzepkowska, Tomasz Jasiński, Małgorzata Domino, Bartosz Pawliński

## Abstract

Equine endometrosis is a degenerative uterine disease characterized by periglandular fibrosis and represents a major cause of subfertility in mares. While stromal cell activation results in pathological tissue remodeling, the involvement of cell-surface integrators (*THY1*) and growth factor receptors (*PDGFRA*) was not researched. This study evaluated the relative mRNA expression of *THY1* and *PDGFRA* across Kenney and Doig (KD) endometrosis categories and specific histopathological features. Additionally, transcriptomic correlation profiles between target genes and markers of myofibroblast differentiation (*ACTA2*), cellular proliferation (*MKI67*), and steroid hormone receptors (*ESR1, PGR*) were analyzed. Endometrial samples (*n* = 47) were quantified using RT-qPCR. Relative mRNA expression of *THY1* differed significantly across KD categories (*p* = 0.014), exhibiting a transient upregulation in category IIA compared with category I (*p* = 0.009) and category IIB (*p* = 0.002). Furthermore, *THY1* relative mRNA expression was significantly elevated in tissues with glandular basal lamina disruption (*p* = 0.022). Conversely, *PDGFRA* relative mRNA expression remained stable across disease categories (*p* > 0.05). *THY1* relative mRNA expression correlated very strongly with *ACTA2* (ρ = 0.875, *p* < 0.001) and strongly with *ESR1* (ρ = 0.652, *p* < 0.001). *PDGFRA* relative mRNA expression exhibited a very strong positive correlation with *ESR1* (ρ = 0.864, *p* < 0.001) and a strong positive correlation with *MKI67* (ρ = 0.674, *p* < 0.001). These results indicate that *THY1* transcription is transiently induced during early stromal remodeling and basal lamina loss, whereas *PDGFRA* maintains constitutive transcript stability. Observed co-expression profiles reflect statistical co-variation among stromal remodeling, proliferative, and steroid-responsive cell populations in bulk tissue.

## 1. Introduction

Equine endometrosis is a major cause of subfertility in mares and a principal constraint on reproductive efficiency in horse breeding worldwide [1,2]. Pathologically, established periglandular collagen deposition impairs endometrial histotroph secretion and placental development, thereby predisposing affected mares to early embryonic death and reduced foaling rates [3,4]. Histopathological grading according to the Kenney and Doig (KD) system categorizes disease severity into categories I, IIA, IIB, and III based on the extent of glandular involvement and periglandular fibrotic layers [2,5]. However, advanced endometrosis represents an irreversible structural state with limited therapeutic options, underscoring the necessity to elucidate the early molecular triggers driving stromal remodeling and tissue degeneration [1,6].

The pathogenesis of endometrial fibrosis is orchestrated by fibroblast activation and their phenotypic transition into contractile myofibroblasts, characterized by upregulated alpha-smooth muscle actin (α-SMA), encoded by *ACTA2* [7,8]. Transforming growth factor beta 1 (TGF-β1), encoded by *TGFB1*, serves as a master profibrotic cytokine driving extracellular matrix (ECM) synthesis, collagen deposition, and myofibroblast differentiation [8–10]. Furthermore, destructive endometrosis is distinguished by focal disruption of the glandular basal lamina, exposing basement membrane components (e.g., laminin and fibronectin) to the surrounding stroma and accelerating epithelial-stromal disorganization [11,12]. These structural disruptions occur within a cyclically regulated hormonal environment, where altered expression of estrogen receptor α (*ESR1*) and progesterone receptor (*PGR*) impairs cyclic responsiveness and stromal-epithelial crosstalk [2,11,13]. Nevertheless, whether early tissue remodeling is governed solely by classical profibrotic signaling (*TGFB1, ACTA2*) or involves specific cell-surface regulators and growth factor receptors remains to be clarified.

Thymus cell antigen 1 (Thy1, CD90), encoded by *THY1*, is a cell-surface glycoprotein that functions as a fundamental integrator of cell-matrix and cell-cell communication [14,15]. Although lacking a direct classical receptor, Thy1 interacts dynamically with integrins (e.g., αvβ3) and proteoglycans within the ECM, regulating cell adhesion, migration, mechanotransduction, and cell survival [14,16]. Importantly, Thy1-integrin interactions mediate the conformational activation of latent TGF-β1 complexes embedded in the extracellular scaffold [9,17]. Moreover, Thy1 is enriched on extracellular vesicles released by activated stromal cells, facilitating intercellular signaling and tissue repair [18]. In various fibrotic pathologies, Thy1 regulates fibroblast differentiation, myofibroblast apoptosis, and tissue regeneration [15,17,18]. In ovine models of induced endometrial fibrosis, upregulation of *THY1* accompanies enhanced cell migration and ECM disorganization [19]. However, the specific expression pattern of *THY1* during equine endometrosis remains unexplored.

Platelet-derived growth factor receptor alpha (PDGFRα), encoded by *PDGFRA*, is a receptor tyrosine kinase primarily expressed on cells of mesenchymal origin, including resident stromal fibroblasts [20,21]. Ligand-induced dimerization of PDGFRα triggers downstream phosphoinositide 3-kinase (PI3K) and extracellular signal-regulated kinase (ERK) cascades, promoting stromal fibroblast survival, cell cycle progression, migration, and ECM synthesis [20–22]. In multi-organ fibrotic conditions (e.g., pulmonary, hepatic, and cardiac fibrosis), PDGFRα activation expands the resident fibroblast pool and prevents myofibroblast apoptosis, sustaining active ECM deposition [23–25]. Furthermore, crosstalk between PDGFRα and non-genomic estrogen receptor α (ERα) signaling has been demonstrated in hormone-responsive tissues, where estrogen can modulate *PDGFRA* transcription and ligand sensitivity [26–28]. In the mare uterus, PDGFRα signaling may contribute to stromal cell homeostasis, hyaluronan synthesis, and matrix metalloproteinase modulation [29–31].

Accordingly, the relative mRNA expression of *THY1* and *PDGFRA* was evaluated in equine endometrial tissue across categories of endometrosis and in relation to defined histopathological features. Furthermore, transcriptomic correlation profiles between *THY1, PDGFRA*, and selected markers of myofibroblast differentiation (*ACTA2*), cellular proliferation (*MKI67*), and steroid hormone receptors (*ESR1* and *PGR*) were analyzed to determine patterns of transcript co-variation during endometrial tissue remodeling.

## 2. Materials and methods

### 2.1 Tissue collection

Post-mortem endometrial tissue specimens (n = 72) were collected from mares of diverse breeds and ages at a commercial slaughterhouse (Rawicz, Poland) in August 2023. As the tissues were collected as by-products of routine commercial slaughter, no procedures were conducted on live animals. This cohort corresponds to the sample set described in our previous study [32]. To avoid anatomical location-induced variability, two adjacent tissue sections were collected from a standardized site within the uterine body, approximately 5 cm from the cervix, from each animal. The first specimen—a full-thickness uterine wall fragment (∼2 cm)—was immediately immersed in 4% buffered formaldehyde (Chempur, Piekary Śląskie, Poland) for subsequent histopathological evaluation. The second specimen, comprising isolated endometrium (∼2 g), was carefully dissected to exclude myometrial contamination. This sample was transferred into RNase-free cryogenic vials (Biologix Group Ltd., Jinan, China), immediately snap-frozen in liquid nitrogen, and stored at –80 °C pending molecular analysis.

Determination of the estrous cycle phase was based on macroscopic ovarian morphology as described by King (2011)[33]. Specifically, the presence of a functional corpus luteum or corpus hemorrhagicum indicated the luteal phase, while the follicular phase was identified by a dominant follicle exceeding 20 mm in diameter without concurrent luteal structures.

### 2.2 Histological processing and Kenney and Doig classification

Formalin-fixed uterine fragments were processed via a standard automated paraffin embedding workflow. Following fixation (24-48 h at room temperature), tissues containing both endometrium and the adjacent myometrial layer were trimmed, placed in cassettes (Simport Scientific, Beloeil, QC, Canada), and dehydrated through an ascending ethanol series (70% to 100%; Avantor Performance Materials Poland S.A., Gliwice, Poland). Specimens were subsequently cleared in xylene (Chempur, Piekary Śląskie, Poland) and infiltrated with paraffin wax (Sigma-Aldrich, St. Louis, MO, USA). Serial sections were cut at a thickness of 4 µm using a rotary microtome and mounted on glass slides (Superfrost, Thermo Fisher Scientific, Waltham, MA, USA).

For morphological evaluation, sections underwent routine hematoxylin and eosin (H&E) staining. Slides were deparaffinized in xylene, rehydrated through descending alcohol grades to distilled water, stained with Harris hematoxylin (Chempur), differentiated in acid alcohol, and counterstained with Eosin Y (Chempur). After final dehydration and clearing, slides were coverslipped using DPX mounting medium (Sigma-Aldrich). Histopathological examination was conducted via light microscopy (BX43, Olympus Corporation, Tokyo, Japan) at multiple magnifications (40× to 400×). Staging was performed by two independent, experienced observers (Ł.Z. and M.D.) who were blinded to both the molecular analysis data and each other’s primary readouts. Any grading discrepancies were resolved via consensus review at a multi-discussion session utilizing the same optical setup. All samples (n = 72) were classified according to the Kenney and Doig (KD) system [5,34]. To eliminate interpretation bias, KD categorization was completed for the entire cohort prior to selecting samples for downstream transcriptomic analysis.

### 2.3 Selection of samples and definition of experimental groups

To investigate endometrosis-associated remodeling while minimizing confounding signals from acute or severe endometritis, a subset of samples (n = 47) was selected for RT-qPCR analysis based on a rigorous histological screening workflow. Samples displaying severe active inflammation, extensive sectioning artifacts, or advanced post-mortem autolysis were excluded from molecular processing.

To capture early-stage or focal fibrotic events that do not meet the strict threshold for category KD IIA at the whole-section level, an additional “early/minor endometrosis” group (designated as I+) was established a priori. The I+ category was operationalized based on the scoring criteria detailed in our previous study [32] and met the following parameters: (i) an overall architectural grading consistent with category KD I; (ii) the concurrent presence of focal remodeling, defined by the presence of active periglandular fibrotic foci or mild glandular degeneration (score 1); (iii) the absence of significant inflammatory infiltration, defined as an overall inflammation score ≤ 2 and a neutrophil infiltration intensity ≤ 1 on the previously established ordinal scales [32].

Samples that were categorized as KD I but did not exhibit these early focal changes were retained as the true histologically normal control group. Accordingly, the final molecular dataset comprised five distinct analytical groups: KD I control (histologically normal endometrium without endometrosis and without endometritis; n = 10), I+ (early/minor endometrosis; n = 7), KD IIA (n = 11), KD IIB (n = 9), and KD III (n = 10).

### 2.4 Additional histopathological assessment

Beyond the standard KD categorization, all endometrial specimens allocated for the molecular dataset underwent evaluation using a structured panel of additional histopathological features. This panel comprised: degeneration of endometrial glands (scale 0-2), periglandular fibrotic foci (0-1) fibroblast activity in fibrotic foci (0-1), basal lamina disruption in endometrial glands (0-1), steroid hormone-dependent glandular differentiation (0-1), luminal epithelium degeneration (0-1), luminal epithelium erosion (0-1), luminal epithelium height (µm), vascular dilation of arterioles and venules (0-1), perivascular fibrosis and remodeling (0-1), overall inflammatory infiltration intensity (0-3), lymphocyte infiltration intensity (0-3), neutrophil infiltration intensity (0-3), macrophage infiltration intensity (0-3), and eosinophil infiltration intensity (0-3). While the precise operational definitions and semi-quantitative scoring scales for each parameter were adopted from our previous study [32], the evaluation protocol was executed as follows. Assessments were performed on H&E-stained sections using a consistent optical configuration (BX43, Olympus Corporation, Tokyo, Japan). Slides were initially screened at low magnification (40× and 100×) to identify well-preserved, representative tissue areas and to exclude regions presenting technical artifacts (e.g., folds, compression, tears) or advanced post-mortem autolysis. Detailed scoring was subsequently conducted at 200× and 400× magnification. For dichotomous (binary) variables, a feature was recorded as present (score 1) if unequivocally identified in at least one representative region of the endometrium; otherwise, it was classified as absent (score 0). For ordinal metrics, the assigned score reflected the predominant morphological pattern across the entire section rather than a localized maximum, unless a focal lesion was sufficiently extensive to alter the overall tissue profile. Stromal inflammatory cell infiltration was scored across five non-overlapping high-power fields (HPFs) per section at a total magnification of 400× (objective 40×, eyepiece 10×). These HPFs were distributed within representative zones of the *stratum compactum* and *stratum spongiosum*. Fields overlapping with glandular or vascular lumina, or containing luminal cellular debris, were strictly avoided to prevent systematic bias in the cellular density estimation. Luminal epithelial height was determined via calibrated digital morphometry using cellSens Standard software (Olympus Corporation). Measurements were taken at a fixed magnification of 400× in intact regions of the luminal epithelium, intentionally excluding eroded areas and gland openings. For each specimen, three distinct measurement points were recorded in µm and averaged to provide a single representative value. All additional parameters were evaluated independently by two investigators (Ł.Z. and M.D.) blinded to the transcriptomic data and to each other’s scores during the primary reading. Discrepancies in feature scoring were resolved through joint consensus sessions using the same microscope configuration (BX43, Olympus Corporation). Categorization thresholds and cell-type identifications adhered to established diagnostic criteria for equine endometrial inflammatory patterns [35,36].

### 2.5 RT-qPCR

Total RNA was isolated from frozen endometrial tissue using the phenol–guanidinium thiocyanate method. Briefly, approximately 100 mg of endometrium per sample was transferred into RNase-free, screw-cap homogenization tubes with ceramic beads (MagNA Lyser Green Beads, Roche Diagnostics, Basel, Switzerland). Tissue was immediately lysed in TRI Reagent (Sigma-Aldrich, St. Louis, MO, USA) and homogenized using a multi-tube bead shaker (MagNA Lyser, Roche Diagnostics, Basel, Switzerland) under standardized settings (frequency: 50-60 Hz, speed 7000 rpm, duration: 20 s; number of cycles: 2), with tubes kept cooled between cycles to limit RNA degradation. RNA extraction was performed using TRI Reagent strictly according to the manufacturer’s protocol, including phase separation with chloroform (Chempur, Piekary Śląskie, Poland), RNA precipitation with isopropanol (Chempur, Piekary Śląskie, Poland), and washing with 75% ethanol prepared from molecular biology grade ethanol (Avantor Performance Materials Poland S.A., Gliwice, Poland) and RNase-free water (Chempur, Piekary Śląskie, Poland). RNA samples were stored at -80 °C until RT-qPCR analysis. RNA concentration and purity were assessed by spectrophotometry using a NanoDrop Lite spectrophotometer (Thermo Fisher Scientific, Waltham, MA, USA). For each sample, the A260/280 ratio was recorded. One-step RT-qPCR was performed on a LightCycler 480 II Real-time PCR system (Roche, Basel, Switzerland) using the TaqMan RNA-to-CT 1-Step Kit (Thermo Fisher Scientific, Waltham, MA, USA). Reactions were set up in 96-well plates (Thermo Fisher Scientific, Waltham, MA, USA) sealed with optical films (Thermo Fisher Scientific, Waltham, MA, USA) in a dedicated clean area to minimize contamination. Each sample was analyzed in technical triplicates. The total reaction volume was 18 µL per well. Predesigned equine TaqMan Gene Expression Assays (Thermo Fisher Scientific, Waltham, MA, USA) were used for quantification of the following transcripts: reference genes: Ec03470487_m1 (*SDHA*) and Ec03210916_gH (*GAPDH*); primary target genes: Ec03470737_m1 (*THY1*) and Ec06993305_m1 (*PDGFRA*); target genes for correlation analyses: Ec03468688_m1 (*IGF1*), Ec03468030_m1 (*TGFB1*), Ec07040967_m1 (*ACTA2*), Ec07039069_g1 (*MKI67*), Ec03467534_m1 (*ESR1*), and Ec03469694_s1 (*PGR*). All assays used a FAM reporter dye. Because commercial predesigned TaqMan assays were used, primer and probe sequences and component concentrations are proprietary to the manufacturer and were therefore not reported; assays were applied at the recommended 1× final concentration.

Thermal cycling conditions were as follows: reverse transcription at 48 °C for 15 min, enzyme activation at 95 °C for 10 min, followed by 40 amplification cycles of denaturation at 95 °C for 15 s and annealing/extension at 60 °C for 60 s. No-template controls (NTCs; RNase-free water substituted for RNA) were included for each assay on every plate to monitor reagent contamination.

Quantification cycle (Cq/Ct) values were calculated using the second derivative maximum method in LightCycler 480 software version 1.5.1.62 (Roche, Basel, Switzerland). The relative mRNA expression was calculated using the comparative Ct method (2^-ΔΔCt^). The geometric mean of two reference genes (*SDHA* and *GAPDH*) was used for normalization. The calibrator was defined as the control group (KD category I without histological signs of inflammation), using the mean ΔCt value of that group as the reference for ΔΔCt calculations. Results are presented as fold change in mRNA expression for categories I+, IIA, IIB, and III relative to healthy endometrium (KD I).

### 2.6 Statistical analysis

Statistical analyses were performed using IBM SPSS Statistics version 29 (Predictive Solutions, Warsaw, Poland), and data visualization was executed in Python (Python Software Foundation, Wilmington, DE, USA). The analytical framework comprised three primary objectives: (i) comparisons of the relative mRNA expressions of *THY1* and *PDGFRA* among the endometrosis categories (KD I, I+, IIA, IIB, and III); (ii) comparisons of the relative mRNA expression between groups defined by additional histopathological features; and (iii) correlation analyses among different transcripts as well as between the relative mRNA expression and ordinal histopathological scores.

Distributional assumptions for quantitative variables were assessed using the Shapiro–Wilk test. Because the relative mRNA expression did not meet normality assumptions, nonparametric tests were applied. Differences in the relative mRNA expression among more than two groups were evaluated using the Kruskal–Wallis test. When the omnibus test indicated a significant difference, it was followed by Dunn’s post-hoc test for multiple pairwise comparisons, with p-values adjusted using the Benjamini–Hochberg procedure to control the false discovery rate (FDR). For comparisons involving dichotomous histopathological features, the relative mRNA expression was compared between groups using the Mann–Whitney U test.

Associations between continuous or ordinal variables were evaluated using Spearman’s rank correlation coefficient (ρ). Correlation strength was interpreted according to Akoglu (2018) as weak (|ρ| = 0.10–0.29), moderate (|ρ| = 0.30–0.59), strong (|ρ| = 0.60–0.79), or very strong (|ρ| = 0.80–1.00) [37]. All statistical tests were two-tailed, and statistical significance was defined as an adjusted *p*-value of less than 0.05 (*p* < 0.05).

## 3. Results

### 3.1 Relative mRNA expression across categories of endometrosis

The relative mRNA expression of *THY1* varied significantly across the investigated endometrosis categories (omnibus Kruskal–Wallis test, *p* = 0.014; Figure 1). Subsequent post-hoc pairwise comparisons utilizing the Benjamini–Hochberg step-up procedure revealed significant differences between category IIA and IIB (*p* = 0.002) as well as between category I and IIA (*p* = 0.009), as both values successfully met the adapted significance thresholds determined by the FDR control. No other pairwise differences in *THY1* transcript levels reached the critical thresholds defined by the procedure. The relative mRNA expression of *PDGFRA* did not differ significantly among any of the experimental categories (Figure 1).

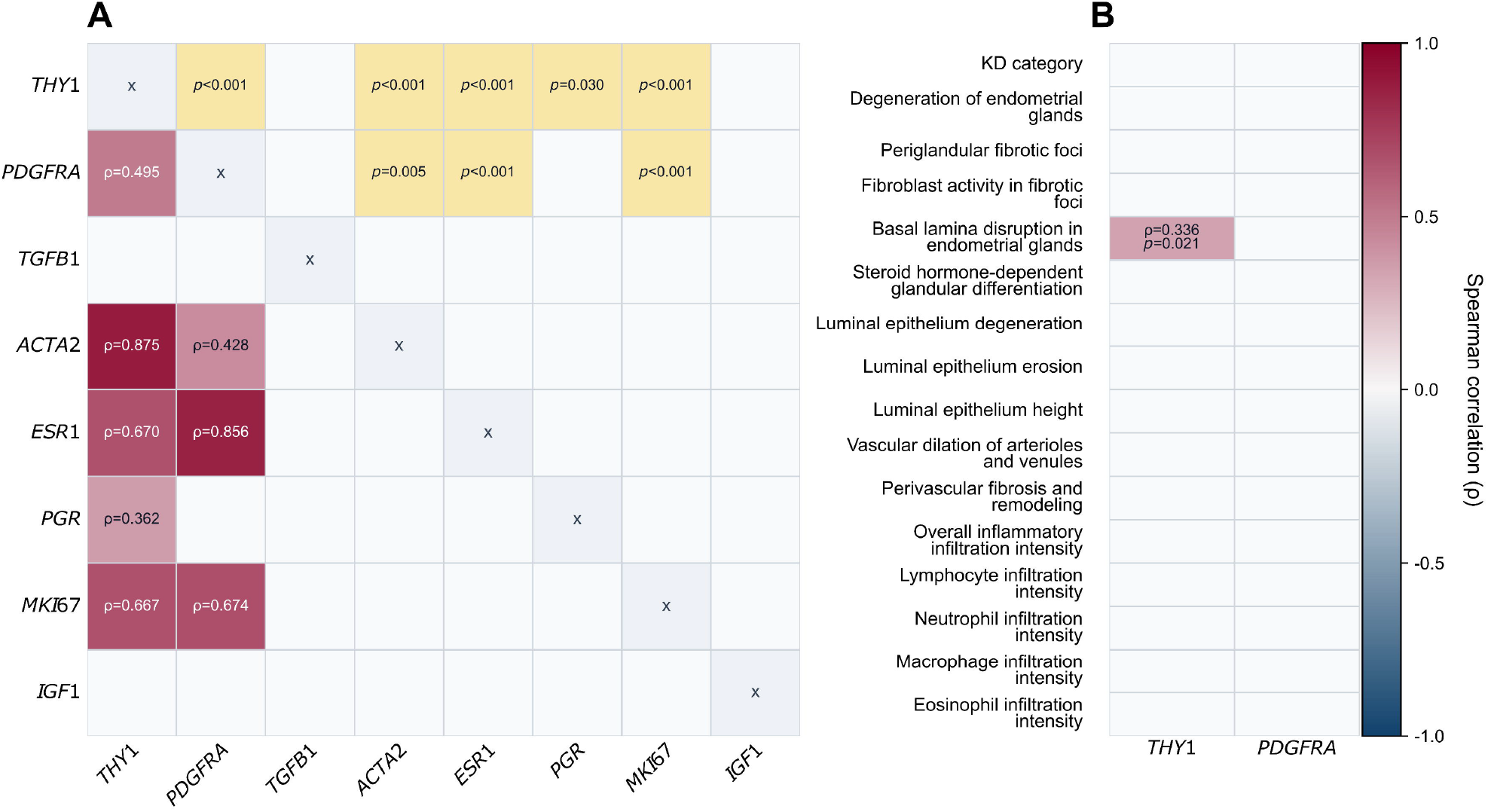

### 3.2 Relative mRNA expression in relation to additional histopathological features

When stratified by specific histopathological features, samples exhibiting basal lamina disruption in the endometrial glands (representative of destructive endometrosis) showed significantly elevated *THY1* mRNA expression compared to nondestructive endometrosis (*p* = 0.022; Figure 2). No other statistical differences reached significance for either *THY1* or *PDGFRA* expression when analyzed against the remaining dichotomous histopathological parameters.

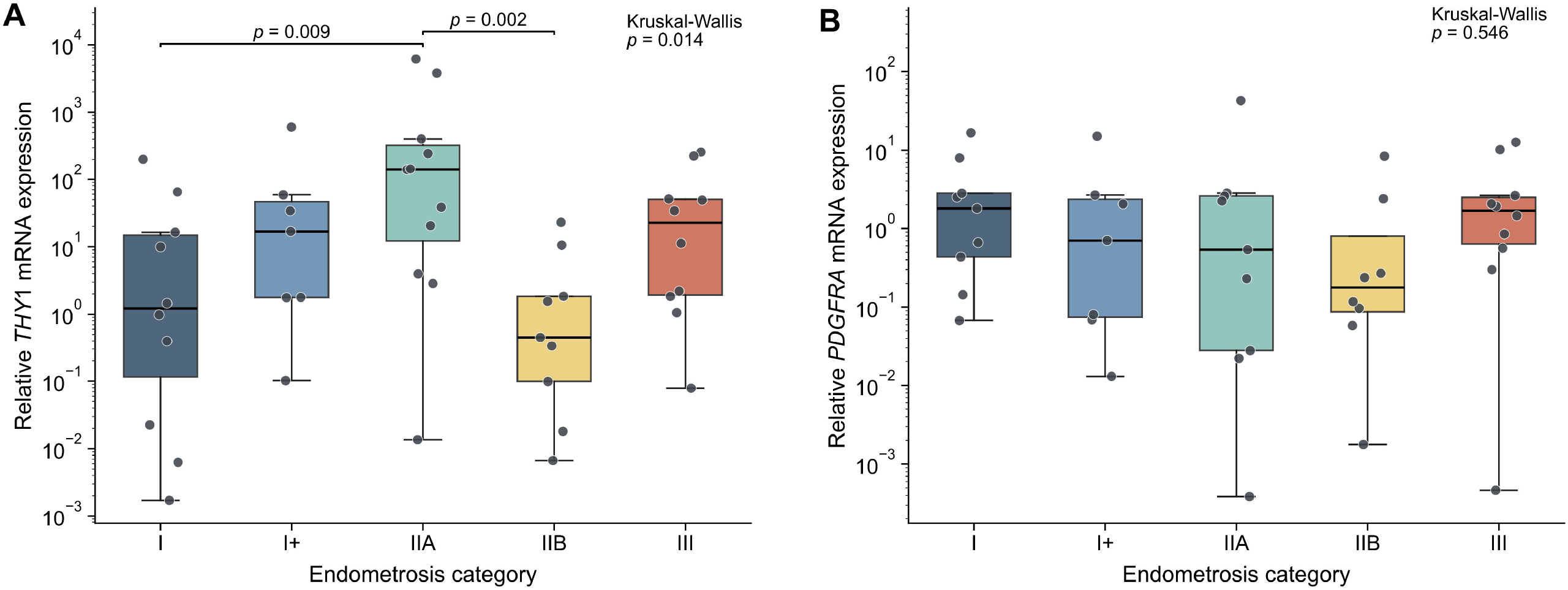

### 3.3 Correlation profiles among mRNA transcripts and histopathological features

The relative mRNA expression of *THY1* correlated very strongly and positively with *ACTA2* (ρ = 0.875, *p* < 0.001), and showed strong positive correlations with *ESR1* (ρ = 0.670, *p* < 0.001), *MKI67* (ρ = 0.667, *p* < 0.001), and moderate positive correlations with *PDGFRA* (ρ = 0.495, *p* < 0.001) and *PGR* (ρ = 0.362, *p* = 0.030). *PDGFRA* expression was positively and very strongly correlated with *ESR1* (ρ = 0.856, *p* < 0.001), strongly correlated with MKI67 (ρ = 0.674, *p* < 0.001) and moderately with *ACTA2* (ρ = 0.428, *p* = 0.005).

In terms of associations between transcripts and morphology, *THY1* expression exhibited a significant positive correlation exclusively with the severity of basal lamina disruption in the endometrial glands (ρ = 0.336, *p* = 0.021). No statistically significant correlations were found between *PDGFRA* mRNA levels and any of the evaluated semi-quantitative histopathological scores (Figure 3).

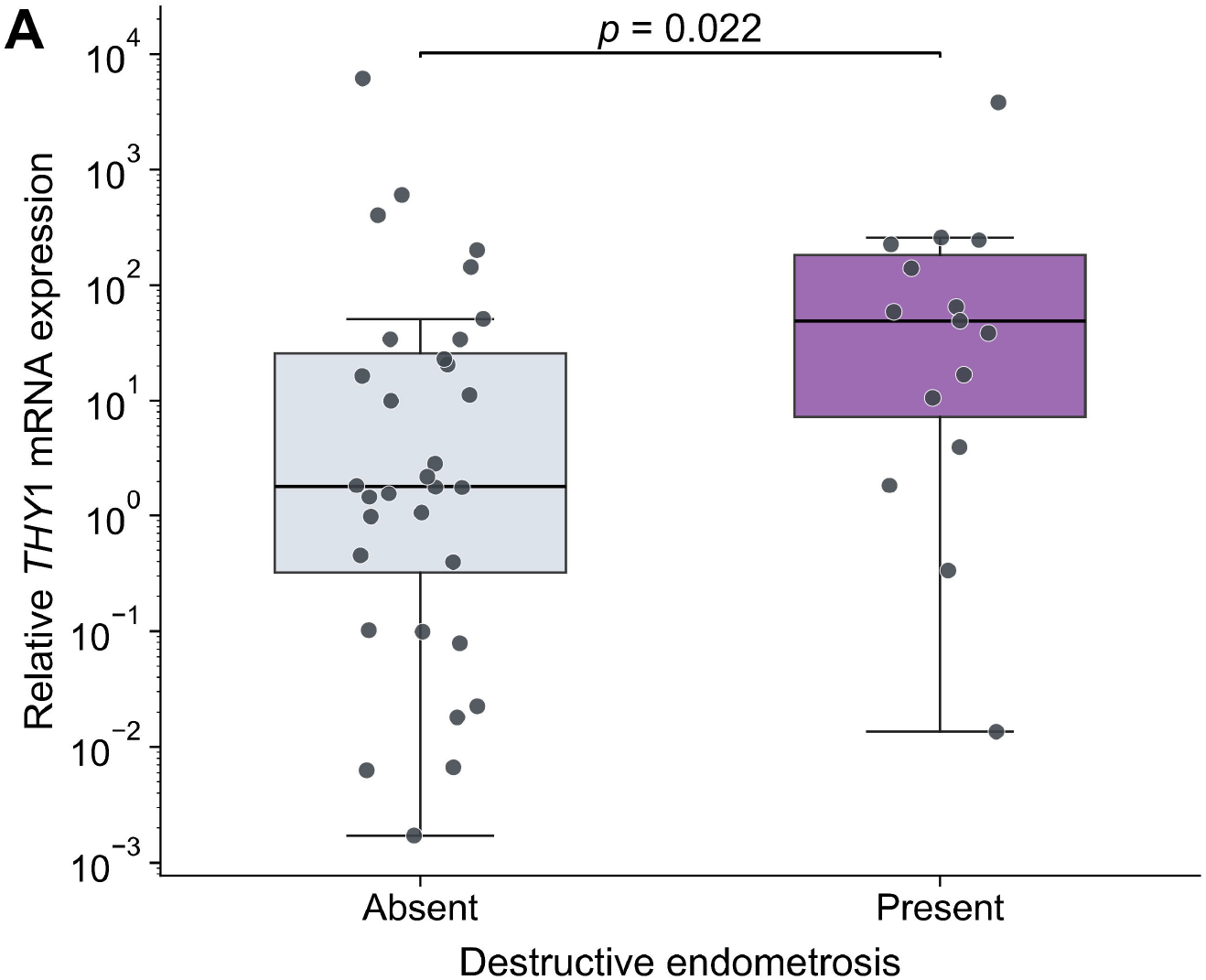

## 4. Discussion

### 4.1 Relative mRNA expression of *THY1* and *PDGFRA* across categories of endometrosis

The relative mRNA expression of *THY1* varied significantly across Kenney and Doig (KD) endometrosis categories, exhibiting a transient upregulation in category IIA (mild endometrosis) relative to category I (normal control) and category IIB (moderate endometrosis), whereas levels in category I+ (early/minor lesions) and categories IIB–III remained comparable to healthy controls. This transient elevation in category IIA implies that Thy1 (CD90) may participate primarily in the early phase of endometrial stromal remodeling rather than driving advanced tissue degeneration. Thy1 is a cell-surface glycoprotein that mediates cell–matrix and cell–cell interactions through binding to integrins and proteoglycans [14]. Integrins are essential for converting latent, inactive transforming growth factor beta 1 (TGF-β1) into its biologically active conformation [9] and modulating membrane receptor crosstalk [14,17]. Given that myofibroblasts exhibit elevated membrane integrin expression during early fibrotic induction [38], the increased transcription of *THY1* in early-stage endometrosis may reflect heightened cellular adhesion, stromal cell migration, and structural reorganization of the extracellular matrix (ECM). Furthermore, altered *THY1* mRNA levels may align with shifts in proteoglycan synthesis within the amorphous ECM, as reported following intra-uterine treatments that induce endometrial fibrosis [39]. Notably, the lack of direct correlation between relative mRNA expression of *TGFB1* and *THY1* suggests that Thy1 and TGF-β1 operate through distinct or non-linear transcriptional regulatory mechanisms during the onset of equine endometrosis.

In contrast to *THY1*, relative mRNA expression of *PDGFRA* did not differ significantly across KD endometrosis categories, maintaining stable baseline expression throughout disease progression. This stability suggests that platelet-derived growth factor receptor alpha (PDGFRα) maintains a constitutive, basal transcript presence within the endometrial stroma. Although, participation of PDGF/PDGFRα axis in the fibrotic activation in equine endometrosis cannot be excluded, as it may be regulated via ligand synthesis (PDGF isoforms), proteolytic ligand activation, or receptor dimerization dynamics rather than transcriptional upregulation of the receptor gene itself [21].

### 4.2 Relative mRNA expression in relation to additional histopathological features

Stratification by specific histopathological lesions revealed significantly elevated *THY1* transcript levels in endometrial specimens presenting glandular basal lamina disruption compared with nondestructive endometrosis. Loss of basal lamina integrity exposes basement membrane components to ECM, which enhances interactions between glandular epithelium, fibroblasts, leukocytes and ECM via Thy1 [18,40]. Increased *THY1* transcription in this setting may also reflect active extracellular vesicle-mediated inter-cellular signaling [18] or, alternatively, an endogenous compensatory response aimed at limiting TGF-β1-driven fibrogenesis and restraining unchecked tissue turnover [17]. The absence of correlation between basal lamina breakdown and inflammatory infiltration scores implies that this Thy1-mediated response operates within the scope of physiological resident leukocyte activity rather than acute inflammatory infiltration [2,35].

### 4.3 Correlation among mRNA transcripts and histopathological features

*THY1* relative mRNA expression demonstrated a very strong positive correlation with *ACTA2* (ρ = 0.875, *p* < 0.001), strong positive correlations with *ESR1* (ρ = 0.670, *p* < 0.001), and *MKI67* (ρ = 0.667, *p* < 0.001), and moderate positive correlations with *PDGFRA* (ρ = 0.495, *p* < 0.001), and *PGR* (ρ = 0.362, *p* = 0.030). However, because transcripts were quantified in bulk tissue homogenates, these statistical relationships cannot be localized to a specific cellular subpopulation or definitively ascribed to a single functional signaling axis; consequently, their biological interpretation remains exploratory. Within these constraints, the strong co-expression of *THY1* and *ACTA2* suggests that Thy1 may participate in, or reflect, phenotypic shifts within the endometrial stromal cell compartment. Furthermore, although Thy1 is primarily recognized for its role in inhibiting apoptosis, the moderate positive correlation observed between *THY1* and *MKI67* expression might tentatively reflect a shift toward a less specialized cellular state, which in turn could be associated with an increased proliferative capacity [14,16]. A moderate positive correlation was also identified between *THY1* and *PDGFRA* transcript levels, raising the possibility of a functional interplay between Thy1 and PDGFRα signaling in maintaining the resident fibroblast population. In parallel, the positive associations with steroid receptor genes (*ESR1* and *PGR*) indicate that *THY1* transcription co-varies with hormone-responsive endometrial components [13,41]. Viewed cautiously, the strong positive correlation between *THY1* and *ESR1* could align with a putative homeostatic or protective role of Thy1 during the initial stages of endometrosis, potentially reflecting the preservation of endometrial functionality and cyclic regulation alongside elevated *THY1* transcription. Finally, given that estrogens also stimulate progesterone receptor synthesis [41], the moderate positive correlation observed between *THY1* and *PGR* expression may represent a secondary effect of this broader estrogen– progesterone regulatory crosstalk rather than a direct functional interaction.

*PDGFRA* relative mRNA expression exhibited a very strong positive correlation with *ESR1* (ρ = 0.856, *p* < 0.001), strong positive correlation with *MKI67* (ρ = 0.674, *p* < 0.001), alongside moderate positive correlations with *THY1* (ρ = 0.495, *p* < 0.001) and *ACTA2* (ρ = 0.428, *p* = 0.005). The marked co-variation between *PDGFRA* and *MKI67* indicates that higher receptor transcript levels coincide with increased cellular proliferation in endometrial samples [20,21]. In human stromal cells and vascular myocytes, PDGFRα activation promotes cell cycle progression [22,42]; however, in bulk endometrial tissue, this correlation may simply reflect the overall abundance of actively proliferating stromal cells rather than direct receptor-driven mitogenesis. Similarly, the moderate positive correlation with *ACTA2* aligns with reports linking PDGFRα presence to stromal cell differentiation and survival [23,24,31]. The very strong correlation between *PDGFRA* and *ESR1* suggests co-expression in hormone-sensitive stromal compartments [28,43]. Although cross-talk between estrogen signaling and PDGFRα pathways has been described in other organs [26,27,44,45], these relationships represent statistical co-occurrences that warrant targeted functional testing.

### 4.4 Limitations of the study

Several methodological limitations inherent to this study warrant explicit consideration to guard against over-interpretation. First, the cross-sectional study design relied on post-mortem tissues collected at a commercial abattoir within a single calendar month. Because these specimens were obtained as routine slaughter by-products, comprehensive clinical metadata—including prior reproductive history, parity, exact age, and systemic health status—were unavailable, precluding longitudinal inferences regarding temporal disease progression. Furthermore, estrous cycle phases were categorized strictly from macroscopic ovarian morphology rather than systemic endocrine profiling (e.g., serum progesterone and estradiol concentrations). Given that steroid receptors (ESR1, PGR) and proliferation markers (MKI67) undergo pronounced cyclic regulation, residual confounding driven by unmeasured hormonal fluctuations cannot be entirely excluded [2,46].

Most importantly, relative mRNA expression was quantified from full-thickness endometrial homogenates using bulk-tissue RT-qPCR. Such homogenates integrate luminal and glandular epithelia, vascular walls, resident stromal fibroblasts, and infiltrating immune cells into a single pooled measurement. Because pathological remodeling—such as periglandular fibrosis, epithelial erosion, or leukocytic influx—alters the proportional representation of these compartments, observed shifts in relative mRNA expression may reflect changes in tissue cellularity rather than altered transcriptional activity within a specific cell subpopulation.

Transcriptomic findings were not corroborated at the protein level via immunohistochemistry, Western blotting, or spatial proteomics. Relative mRNA expression does not invariably correspond to protein abundance, subcellular localization, or functional biological activity. For instance, receptors such as PDGFRα require ligand-induced dimerization and intracellular phosphorylation to initiate downstream profibrotic cascades [21], whereas key mediators like TGF-β1 are secreted as latent complexes requiring proteolytic or integrin-mediated activation [9]. Without compartment-specific protein localization and functional activation assays, mechanistic inferences regarding Thy1/PDGFRα signaling cross-talk or stromal phenotypic transitions remain speculative. Finally, although histopathological evaluation followed standardized criteria, ordinal scoring remains inherently semi-quantitative. Consequently, the associations reported here should be viewed as exploratory, hypothesis-generating signals that require validation in prospective clinical cohorts incorporating spatial and protein-level verification.

## 5. Conclusions

In conclusion, this study demonstrates a transient upregulation of *THY1* transcript abundance in mild equine endometrosis (category IIA) and in endometrial tissues exhibiting glandular basal lamina disruption, whereas *PDGFRA* transcription remains stable across categories. Strong positive correlations were identified between *THY1* and *ACTA2*, as well as between *PDGFRA, MKI67*, and *ESR1*, highlighting significant statistical co-variation among transcripts associated with stromal remodeling, cellular proliferation, and steroid responsiveness in bulk tissue. However, because these findings rely on whole-tissue RT-qPCR without spatial or protein-level functional verification, these co-expression patterns should be cautiously interpreted as parallel observational signals rather than evidence of a single unified regulatory pathway, serving as a hypothesis-generating foundation for future cell-type-specific translational research.

## CRediT authorship contribution statement

**Łukasz Zdrojkowski:**Conceptualization, Formal analysis, Investigation, Data Curation, Writing - Original Draft, Funding acquisition. **Anna Niwi**ń**ska:** Formal analysis, Investigation. **Ewa Kautz:** Methodology, Investigation. **Dawid Tobolski:** Formal analysis, Data Curation, Visualization Writing - Review & Editing. **Magdalena Fajkowska:** Methodology, Validation, Writing - Review & Editing. **Małgorzata Rzepkowska:** Methodology, Validation, Writing - Review & Editing. **Tomasz Jasi**ń**ski:** Validation, Writing - Original Draft, Writing - Review & Editing. **Małgorzata Domino:** Investigation, Validation, Resources, Project administration, Funding acquisition. **Bartosz Pawli**ń**ski:** Conceptualization, Resources, Supervision.

## Data and availability statement

This paper forms part of the PhD Thesis of Łukasz Zdrojkowski (2025). The data generated during this study have been deposited in the Zenodo repository at 10.5281/zenodo.20326518. Further information is available from the corresponding author on reasonable request.

